# Perinatal Inflammation Disturbs Secondary Germinal Zone Neurogenesis and Gliogenesis Producing Deficits in Sociability

**DOI:** 10.1101/2021.08.18.456694

**Authors:** Fernando Janczur Velloso, Anna Wadhwa, Ekta Kumari, Ioana Carcea, Ozlem Gunal, Steven W. Levison

**Affiliations:** Department of Pharmacology, Physiology & Neuroscience, New Jersey Medical School, Rutgers University, Newark, NJ, USA 07103; Department of Psychiatry, New Jersey Medical School, Rutgers University, Newark, NJ, USA 07103

**Author notes:** **Correspondence should be addressed to:** Steven W. Levison, PhD, Department Pharmacology, Physiology & Neuroscience, New Jersey Medical School, Rutgers University, 205 S. Orange Ave, Newark, NJ 07103. <u>**Author Contributions:** FJV, AW and EK were</u> responsible for performing all of the experiments reported in this manuscript and the data analysis. IC, OG and SWL designed all of the studies. FJV wrote the first draft of the manuscript, and all of the authors thoroughly discussed the results and their interpretation and approved the final manuscript. <u>**Author Disclosure Statement:**</u> The authors declare that they have no relevant or material financial interests that relate to the research described in this paper. <u>**Data Availability:**</u> All data will be made available for inspection or use upon request to the corresponding author.

**Keywords:** subventricular zone, subgranular zone, cytokines, Interleukin-6, cell proliferation, stem cells

## Abstract

Epidemiologic studies have demonstrated that infections during pregnancy increase the risk of offspring developing Schizophrenia, Autism, Depression and Bipolar Disorder and have implicated interleukin-6 (IL-6) as a causal agent. However, other cytokines have been associated with psychiatric disorders; therefore, it remains to be established whether elevating IL-6 is sufficient to alter the trajectory of neural development. Furthermore, most rodent studies have manipulated the maternal immune system at mid-gestation, which affects the stem cells and progenitors in both the primary and secondary germinal matrices. Therefore, a question that remains to be addressed is whether elevating IL-6 when the secondary germinal matrices are most active will affect brain development. Here, we have increased IL-6 from postnatal days 3-6, when the secondary germinal matrices are rapidly expanding. Using Nestin-Cre^ERT2^ fate mapping we show that this transient increase in IL-6 decreased neurogenesis in the dentate gyrus of the dorsal hippocampus, reduced astrogliogenesis in the prefrontal cortex and amygdala and decreased oligodendrogenesis in the body and splenium of the corpus callosum all by ∼50%. Moreover, the IL-6 treatment elicited behavioral changes classically associated with neurodevelopmental disorders. As adults, IL-6 injected male mice lost social preference in the social approach test, spent ∼30% less time socially engaging with sexually receptive females and produced ∼50% fewer ultrasonic vocalizations during mating. They also engaged ∼50% more time in self-grooming behavior and had an increase in inhibitory avoidance. Altogether, these data provide new insights into the biological mechanisms linking perinatal immune activation to complex neurodevelopmental brain disorders.

**Significance statement:** In these studies, we doubled circulating IL-6 levels in mice from postnatal days 3-6 to test the hypothesis that this would be sufficient to disturb neural development. More specifically, we hypothesized that IL-6 would affect postnatal neural stem cell and progenitor expansion and specification. We show that this transient increase in IL-6 decreases the numbers of granule neurons in the dorsal hippocampus, astrocytes in the prefrontal cortex and amygdala and oligodendrocytes in the corpus callosum. Importantly, this transient increase in IL-6 changes the sociability, communication and repetitive behaviors of the treated mice as adults, which are core symptoms pertinent to several neurodevelopmental psychiatric disorders.

## Introduction

Epidemiologic studies have revealed that perinatal infections, autoimmune reactions, allergies, exposure to pesticides and air pollution during pregnancy increase the risk of offspring developing neurodevelopmental disorders that include schizophrenia, autism spectrum disorder (ASD), Tourette disorder, depression, developmental delay and cerebral palsy (Mednick et al., 1988; Zerbo et al., 2015; Bear and Wu, 2016; Carter and Blizard, 2016; Jiang et al., 2016; Han et al., 2021) These risk factors converge on the activation of the maternal immune system, however, the mechanisms linking inflammation to abnormal perinatal brain development are poorly understood.

A number of cytokines and chemokines are produced during inflammation that are consistently associated with neurodevelopmental disorders. Interleukin-6 (IL-6) in particular has been associated with schizophrenia, ASD and major depressive disorder (Garay and McAllister, 2010). A retrospective study that analyzed the mid-gestational serum levels of 415 mothers of children with ASD found that high levels of IL-6 were specifically associated with ASD cases with intellectual disabilities (Jones et al., 2017). Moreover, elevated levels of IL-6 and IL-5 were found in the amniotic fluid of women who’s children were later diagnosed with psychiatric disorders (Abdallah et al., 2013).

Levels of IL-6 also are elevated postnatally in children who develop ASD. In a study that evaluated bloodspots in newborns, children who were subsequently diagnosed with ASD had increased levels of IL-6 and IL-8 compared with controls (Heuer et al., 2019). Furthermore, IL-6 is elevated two-fold in plasma collected from ASD children and levels of IL-6 are directly associated with increased stereotypy (Ashwood et al., 2011). Accordingly, IL-6 is being increasingly considered to be a biomarker for ASD (Yang et al., 2015).

Rodent models of maternal immune activation (MIA) using bacterial lipopolysaccharide (LPS) or viral (Poly(I:C)) mimetics have confirmed that elevating inflammatory cytokines prenatally produces long lasting cognitive and behavioral changes (Shi et al., 2003; Smith et al., 2007). Several studies have shown that IL-6 is a key pathogenetic cytokine (Samuelsson et al., 2006). For instance, Smith et al. (2007) showed that a single injection of IL-6 into pregnant mice at mid-gestation produced the same cognitive and behavioral deficits as poly(I:C). Moreover, antagonizing IL-6 (but not TNFα or IL-1β) was sufficient to attenuate the behavioral and immunological abnormalities induced by poly(I:C) (Smith et al., 2007). Similarly, ASD-like behaviors induced by LPS can be attenuated by controlling plasma IL-6 levels postnatally with the anti-inflammatory drug Pioglitazone (Kirsten et al., 2018). Furthermore, pharmacologically inhibiting IL-1β, IL-6 and TNF-α attenuates stereotyped behaviors and enhances social interactions in the BTBR mice model for autism (Ahmad et al., 2019).

The IL-6 in the maternal serum can cross the placenta and the fetal blood-brain barrier (Banks et al., 1994; Dahlgren et al., 2006), resulting in increased levels of IL-6 in the fetal circulation and CSF (Zaretsky et al., 2004; Aaltonen et al., 2005). Our studies as well as those of others have shown that IL-6 can directly affect the neural stem cells (NSCs) and neural progenitors (NPs) in the secondary germinal matrices of the brain, namely, the subgranular zone (SGZ) and the subventricular zone (SVZ). These progenitors will produce neurons, astrocytes and oligodendrocytes that populate several late developing brain structures that are relevant to the behavioral deficits seen in neurodevelopmental disorders. The SVZ and SGZ rapidly expand during the early 3rd trimester of human development (Monje et al., 2003; Covey et al., 2011; Wei et al., 2011; Storer et al., 2018; Kumari et al., 2020), and in early postnatal development in mice (Semple et al., 2013; Salmaso et al., 2014). While MIA studies conducted to date have manipulated the immune system at mid-gestation, it remains to be established whether elevating IL-6 postnatally in mice is sufficient to alter the trajectory of neural development and elicit behavioral aberrations. To address these gaps in knowledge, we report here the consequences of elevating IL-6 in mice from postnatal day 3 (P3) to P6.

## Methods

### Mice

Swiss Webster timed-pregnant females were purchased from Charles River laboratories and maintained in the Comparative Medical Resources animal facility at Rutgers University Biomedical Health Sciences. All experiments were performed in accordance with the approved Rutgers University IACUC protocol # 999901108 and were in accordance with National Institute of Health Guide for the Care and Use of Laboratory Animals (NIH publication No. 80-23). For *in vivo* studies, offspring at postnatal day 3 to 6 received 7 intraperitoneal injections with either PBS (controls) or 75 ng of carrier free recombinant mouse interleukin-6 (rmIL6, R&D Systems, Minneapolis MN).

### Barnes maze

The Barnes maze was used to assess spatial memory(Patil et al., 2009) in male and female P42 mice, as described in(Veerasammy et al., 2020a). Briefly, after 1-hour acclimation, animals performed 4 daily trials during 4 days of acquisition training phase, in which the animals had 5 min to reach the escape box using visual cues placed around the Barnes maze. On day 5, a probe trial was performed for 90 seconds (escape box closed). At day 12, memory retention was evaluated by repeating the probe task. The Ethovision XT system (Noldus Information Technologies) was used for video-tracking and analysis. All trials performed under 340 lux.

### Elevated plus maze (EPM)

The Elevated plus maze was used to assess anxiety (Walf and Frye, 2007) in male and female mice at P42-P65, as described in (Veerasammy et al., 2020b). Briefly, following a 1-hour habituation (40 lux), mice were placed on the center of the apparatus facing the same open arm and tracked for 5 min using the Ethovision XT system.

### Open field test (OFT)

Locomotion and anxiety were assessed at P65 for both male and female mice using the Open Field Test (OFT). After 1 hour of acclimation, the mice were placed in the center of the arena (60 × 60 × 30 cm, 40 lux) and allowed to explore it for 10 min. The Ethovision XT system was used for tracking and analysis. The inner area was set as consisting of 25% of the total area of the arena).

### Self-grooming

Male and female P65 mice were assessed for spontaneous self-grooming individually in clean standard mouse cages (46×20 cm) under dim light (40 lux). After a 5-minute habituation period, mice were videotaped and a highly trained observer, who remained blind to the experimental group, scored cumulative time spent grooming any region of the body over 10-minute test sessions.

### Social approach and novel social partner

Social approach was assayed in P42 male mice using an automated 3-chamber apparatus, consisting of a 35 × 45 cm arena divided into 3 equal sized chambers with gates to control free movement across chambers(Yang et al., 2011). After 1-hour acclimation in single-housed cages, test animals were allowed to freely explore the 3-chamber apparatus for 5 min (open gates and no stimuli) and were then restricted to the central chamber (gates closed). Wire cylinders (15 cm of diameter) were introduced into the side chambers either empty or containing a strain, sex, and age-matched animal socially unfamiliar to the test subject (previously habituated to the enclosure). Gates were opened to allow the test animal to freely move between chambers for 10 min. Cumulative time spent in the social (unfamiliar animal) versus the non-social (empty) chamber was scored using the Ethovision XT live-tracking system. Sniffing time was assessed as the time the head portion of each animal remained inside the sniffing zone (circle 1.5 the radius of the wire cage). Locomotion parameters (total distance, average speed, and freeze time) were also scored.

The social novel partner test was performed using the same 3-chamber apparatus but for this task, the subject was exposed to one chamber that contained a familiar partner (littermate), while the other chamber contained an unfamiliar (novel) age, sex, and strain-matched mouse. The olfaction test in the 3-chamber apparatus was performed as described above, substituting the animals with filter paper with either mouse urine or clean water (control). Enclosures were covered with dark paper to block animal’s view of the inside.

### Inhibitory avoidance (IA)

P42 mice were tested in the inhibitory avoidance task to assess aversive memory(Atucha and Roozendaal, 2015). The IA chamber (Model ENV-010MC, Med Associates, Fairfax VT) was located in a non-illuminated room and consist of a rectangular-shaped box divided into a safe compartment (white and illuminated) and a shock compartment (black and dark). Foot shocks were delivered through the grid floor of the dark chamber. A 1-hour acclimation period was followed by a training session, in which animals were placed in the safe compartment. After 10 s, the door separating the two compartments was opened, allowing access to the dark compartment, where after 1 s the mice received a brief foot-shock (0.7 mA for 3 s). Memory retention was tested at different time points after training (6, 24 and 48 h). Acquisition was measured, in seconds, as the latency to enter the shock compartment.

### Complex horizontal ladder

Adult mice (>P60) were tested using a complex horizontal ladder to assess fine locomotor skills(Antonow-Schlorke et al., 2013). The ladder consisted of two parallel plexiglas walls with transversal metal rods (0.5 in in diameter, 1 cm apart. Mice were tasked with traversing the ladder to reach a dark box. Subjects had 3 training trials one day prior to testing. At least 3 complete crossings were recorded during testing sessions. Trained observers scored the number foot-slips and missteps.

### Hot plate

Tactile pain thresholds were determined for adult mice (>P60) on the hot plate task(Bannon and Malmberg, 2007). After 1h acclimation, the mice were placed on the surface (30cm × 30cm) of a digitally controlled hot plate (model I-39, Campden Instruments, Loughborough, England), heated to a maximum temperature of 55°C. A polypropylene cylinder prevented free roaming during trials. A trained observer scored the latency for the initial pain-related response (licking or flicking the hind paw or jumping). Mice were immediately removed from the surface once a thermal response was observed or after 30 s, to prevent injury.

### Mating behaviors

Male mating behaviors were assessed at P60 as a measure of social interaction. In short, sexually naïve male mice were introduced to an age-matched receptive female partner in a neutral environment and allowed to interact for 16 h before being returned to their home cage. After 48h, males and females were moved to the procedure room but kept isolated for a 1-hour acclimation. Males were placed into a clean cage that had fresh bedding and allowed to habituate for 15 min, after which the female was introduced, and partners freely interacted for 5 min. Sessions were videotaped and scored for mating behaviors by a trained observer blinded to the experimental groups. Behaviors included nose-to-nose sniffing (cumulative time), urogenital sniffing (cumulative time), mounting attempts (frequency) and latency for first mount.

### Ultrasonic vocalization calls

Ultrasonic vocalizations (USV) were recorded during mating sessions. USV calls were recorded using an overhead mounted (30 cm high) high-quality condenser microphone, connected to an Avisoft-UltraSoundGate amplifier (116H). Acoustic data were recorded with a sampling rate of 250,000 Hz in 16-bit format in Avisoft RECORDER and analyzed in SASlab pro software (Avisoft Bioacustics, Nordbahn, Germany). Parameters measured included: Number of ultrasonic calls, interval between calls, average frequency, bandwidth, amplitude, and duration of calls. USV syllable morphology classification was performed in Mouse Ultrasonic Profile ExTraction (MUPET)(Van Segbroeck et al., 2017).

### Fate mapping

To establish the fates of cells arising from SVZ progenitors and stem cells we employed a Nestin-CreERT2/Rosa26-TdTomato reporter mice line. Nestin CreERT2 -/+ line C57BL/6-Tg(Nes-cre/ERT2)KEisc(Lagace et al., 2007) mice were crossed with the TDT +/+ reporter mice B6.Cg-Gt(ROSA)26Sortm14(CAG-tdTomato)Hze/J (Jax mouse stock # 007905)(Madisen et al., 2010). For lineage tracing, only Nestin CreERT2 ^-/+^ TDT ^+/+^ mice were used. Cre recombinase was induced in the stem cells by a single subcutaneous Tamoxifen (75 mg/kg dissolved in a corn oil and EtOH [9:1]; Sigma T5648) injection at P1. Tamoxifen injected mice received IL-6 or PBS injections and were terminated at P36 (**Fig.1**). Brains were sectioned at 30 µm and stained for NeuN (Millipore, Billerica, MA, MAB377, 1/200), S100b (Sigma, St. Louis, MO, S2532, 1/500) and Olig2 (Millipore AB9610, 1/250). Cell numbers were quantified by averaging at least 10 images obtained at 10X. Animals from at least 3 different litters were analyzed for each condition to avoid litter effects.

**Fig.1.**
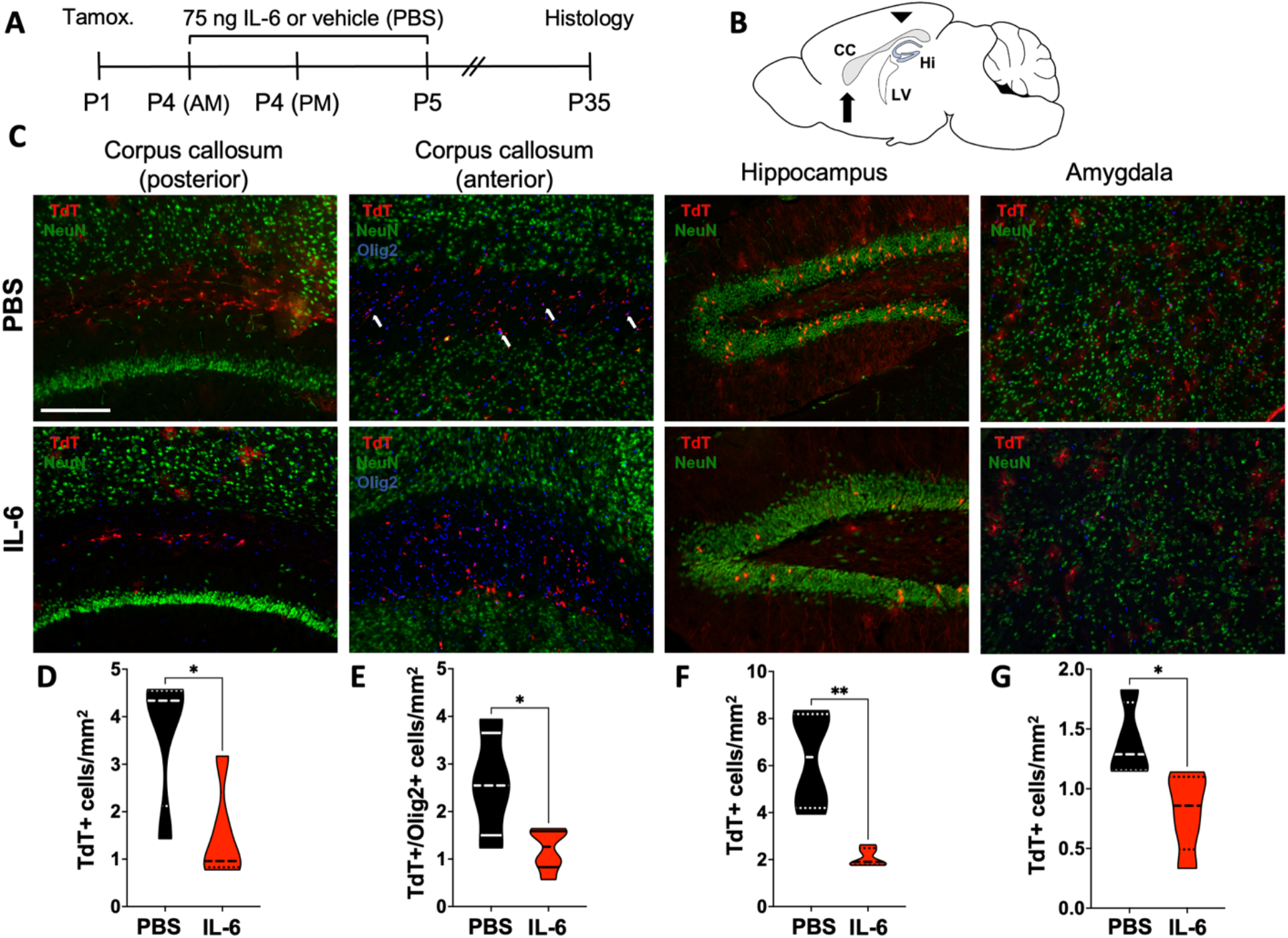
IL-6-treated mice have reduced gliogenesis in the corpus callosum and amygdala and reduced neurogenesis in the hippocampus. The developmental fates of late progenitors were tracked using Nestin-CreERT2/Rosa26-TdTomato reporter mice that had been treated with IL-6 or PBS (control). **A**. Experimental design. Animals received one Tamoxifen injection at P1, two injections of either PBS or IL-6 at P4 (morning [AM] and afternoon [PM]) and third injection (PBS/IL-6) at P5 (morning) **B**. TdTomato+ cell numbers were assessed in anterior (arrow) and posterior (arrowhead) regions of the corpus callosum (CC). LV=Lateral ventricle, Hi=Hippocampus. (**C**) Fluorescence for TdTomato (red), NeuN (green) and Olig2 (blue) positive cells in posterior and anterior regions of the Corpus callosum, in the dentate gyrus of the Hippocampus and in the Amygdala. Blue labelling in the posterior corpus callosum is DAPI. (**D,E,F,G**) Violin plots show quantification of the number of TdTomato+ and TdTomato+/Olig2+ double positive cells per mm^2^ (indicated by white arrows in **C**). Scale bar = 200 µm. *p<0.05, **p < 0.01, by Student’s t-test, n=5 mice/group

### Statistical analysis

Power analyses were performed to estimate the group size required for each experiment based on pilot studies. For *in vivo* studies, at least 8 animals from 3 to 6 litters, were examined per group experimental, at each time point. Animals from one litter were always equally split between groups. Statistical analyses were performed using GraphPad Prism 9 software (GraphPad Software, San Diego, CA). Graphs were produced in Graphpad 9 as violin plots, to express the complexity of the data. Results were analyzed for statistical significance using a two-tailed, unpaired Student’s t-test, one-way or two-way ANOVA followed by Bonferroni’s test for multiple comparison, according to number of means and conditions. Comparisons were interpreted as significant when associated with p < 0.05. Litters were also analyzed individually for all statistical tests to exclude underlying litter effects.

## Results

### Transiently elevating circulating IL-6 reduces gliogenesis in the corpus callosum and amygdala, and neurogenesis in the hippocampus

We previously showed that transiently increasing systemic levels of IL-6 in neonatal mice affects the balance of stem cells and progenitors within the SVZ and an earlier study had shown that increased circulating levels of IL-6 affects neurogenesis in the hippocampus (Monje et al., 2003; Kumari et al., 2020). Therefore, we postulated that neurogenesis and gliogenesis would be abnormal in IL-6 injected (Rx) mice. To test this hypothesis, we used an inducible Nestin-CreERT2/Rosa26-TdTomato reporter mouse line (Lagace et al., 2007) to follow the fates of the stem cells and progenitors in several late developing brain regions. Reporter mice received one Tamoxifen injection at P1 to induce TdTomato expression in the stem cells and then received 75 ng of IL-6 or PBS (vehicle) twice on P4 and once on P5. TdTomato expressing cells were evaluated at P35 (**Fig 1A**). As the subventricular zone is the largest reservoir of progenitors that produce the oligodendrocytes of the white matter, we evaluated gliogenesis with two regions of the corpus callosum. In the more posterior region of the corpus callosum (dorsal to the hippocampus; **Fig.1B)**, there were ∼75% fewer TdTomato+ cells in the IL-6 Rx mice compared to the PBS Rx controls (**Fig.1C,D**; PBS 3.67±0.75 vs. IL-6 1.49±0.45; t(7) = 2.62, p = 0.034, unpaired t-test). There was a similar decrease in TdTomato+ cells in a more anterior region of the corpus callosum of the IL-6 Rx mice (**Fig.1C**; PBS 2.57 ± 0.56 vs. IL-6 1.2 ± 0.18, p < 0.05). The TdTomato+ cells in both regions of the corpus callosum were predominantly small round cells with few processes, suggesting that they were oligodendrocytes. We evaluated the identity of these TdTomato+ oligodendrocytes by co-labelling with the oligodendrocyte lineage marker Olig2 which showed that virtually all of the TdTomato+ cells in the white matter expressed Olig2 (**Fig.2A**). Therefore, we quantified the percentage of TdTomato+/Olig2+ double-positive cells in the anterior corpus callosum which showed that the IL-6 Rx mice had ∼50% fewer postnatally derived oligodendrocytes (**Fig.1E**; PBS 2.57±0.56 vs. IL-6 1.2±0.18; t(8) = 2.75, p = 0.025, unpaired t-test).

**Fig.2.**
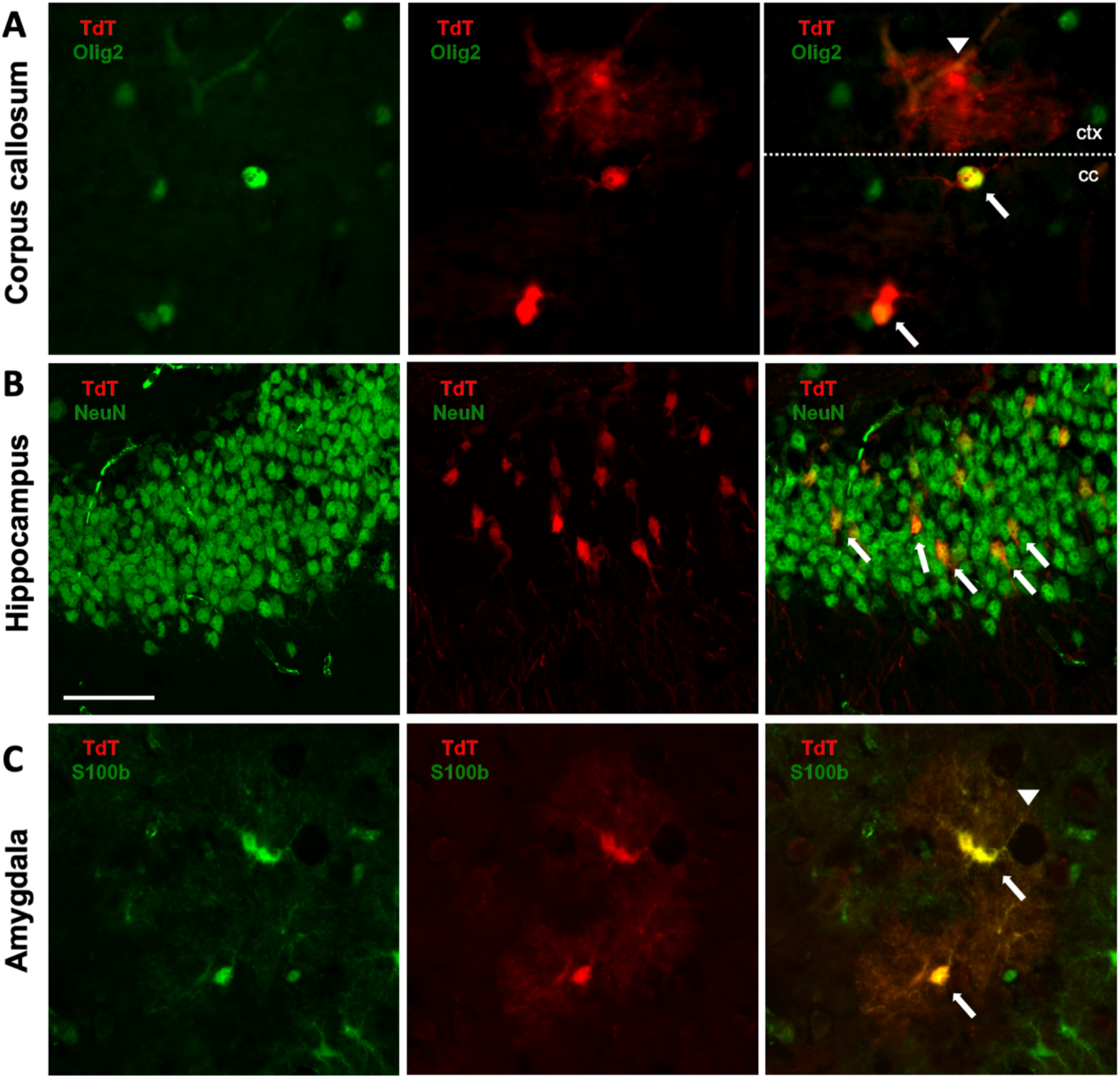
TdTomato+ progenitors give rise to oligodendrocytes in the corpus callosum, neurons in the dentate gyrus of the hippocampus and astrocytes in the amygdala. Late progenitors produced distinct cell types in different regions of the CNS of the Nestin-CreERT2/Rosa26-TdTomato reporter mice. TdTomato co-localized with the oligodendrocyte lineage marker Olig2 in the corpus callosum (**A**), the neuronal marker NeuN in the hippocampus (**B**) and with the astrocyte marker S100b in the amygdala (**C**). The white arrows indicate dual labelled cells. Arrowhead in **A** indicates a neighboring neocortical (ctx) TdTomato+/Olig2-astrocyte at the border of the corpus callosum (cc) to illustrate the morphological differences from the TdTomato+/Olig2+ oligodendrocyte. Arrowhead in **C** marks a blood vessel. Scale bar = 50 µm.

We also evaluated two structures whose function is relevant to social interactions and aversive memory: the hippocampus and the amygdala. Within the hippocampus, the majority of the TdTomato+ cells were round, NeuN+ granule neurons located within the dentate gyrus (**Fig.1C,2B**). IL-6 Rx mice produced ∼70% fewer TdTomato+ neurons in the dentate gyrus compared to control littermates (**Fig.1F**; PBS 6.25±1.06 vs. IL-6 2.09±0.17; t(7) = 4.36, p = 0.003, unpaired t-test). In the amygdala, most of the cells expressing TdTomato were S-100b positive, consistent with their identity as protoplasmic astrocytes. These cells had three to four thick processes that tapered and branched as they extended away from the cell body and some of these cells had processes that wrapped around blood vessels (**Fig.1C,2C**). IL-6 Rx mice produced ∼50% fewer TdTomato+ astrocytes in the amygdala compared to control littermates (**Fig.1G**; PBS 1.39 ±0.16 vs. IL-6 0.81 ± 0.15; t(7) = 2.68, p = 0.031, unpaired t-test)

### IL-6 Rx mice have sex dependent deficits in the Barnes maze, not associated with increased anxiety

As the offspring of dams whose immune systems had been activated during pregnancy display cognitive deficits (Murray et al., 2017), we evaluated the IL-6 Rx mice using the Barnes maze, which is used to assess spatial memory over successive trials (**Fig.3A**). At 6 weeks of age the IL-6 Rx mice of both sexes had a progressively worsening deficit in the latency to complete the maze starting from the second day of training acquisition phase (Two-way ANOVA, main effect for treatment in males F(1,74) = 10.11, p = 0.002, and females F(1,41) = 122.9, p<0.0001, n=8 per group from 3 litters; **Fig.3B**). During the four days of training, mice were tasked with finding the correct escape hole out of 16 possible exits. Failure to improve exit times over this period is usually indicative of a spatial memory impairment. However, while the latencies increased during training for IL-6 Rx mice, especially for males, this was more associated with longer freezing times, rather than errors during exploration. Interestingly, this effect was more pronounced in the females during the 4 days of training, which were significantly more prone to freezing in the IL-6 Rx group (average freezing time PBS 22.72±1.17 vs IL-6 36.48±1.51 F(1,46) = 39.54, p<0.0001, ANOVA, n=8). Furthermore, these changes were not sustained in the male mice during the probe and memory retention phases of the Barnes maze (**Fig 3.A,C,D**). Interestingly, the IL-6 Rx female mice also showed an increased latency to find the escape hole during the training trials, and this persisted through the probe trial and into the retention trial (PBS 7.35±2.55 vs IL-6 45.05±11.7; t(12) = 3.15, p = 0.0084, unpaired t-test). The females had ∼100% longer freeze times (Student’s t-test, PBS 14.69±3.35 vs IL-6 32.36±6.9, t(12) = 2.3, p = 0.04, unpaired t-test) but they made significantly fewer errors in both the probe trial and the retention trials over their PBS Rx littermates (PBS 46±5.73 vs IL-6 26.71±4.16, t(12) = 2.72, p = 0.018 unpaired t-test; **Fig.3C-D**).

**Fig.3.**
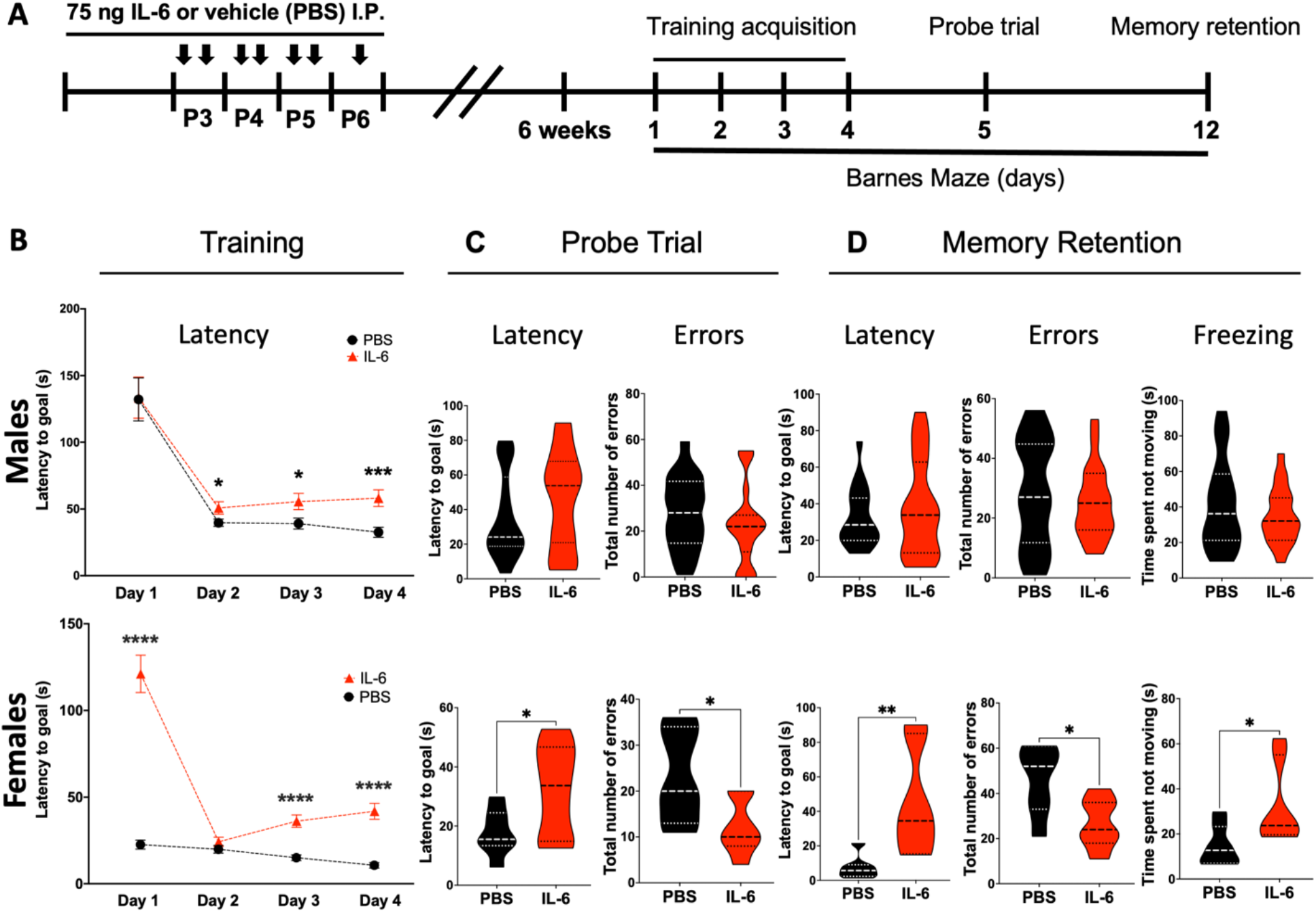
Animals exposed postnatally to IL-6 have sex dependent deficits in the Barnes maze. IL-6 Rx male and female mice were evaluated in the Barnes maze at P42. **A**. Experimental design for twice-daily injections of either 75ng of IL-6 or PBS (control) and the Barnes maze trials. **B**. Latency (s) to find the goal (escape box) during the 4 days of acquisition training. **C**. Latency to find the goal and total errors during the probe trial. **D**. Latency, errors and freezing (percentual of the time the animal was not moving) during memory retention. * p<0.05, ** p<0.01, *** p <0.001, **** p <0.0001 by Student’s t-test (probe and retention) or two-way ANOVA followed by Bonferroni post-hoc test for multiple comparisons (training), n=8 mice per group from 3 litters.

To determine whether the deficits observed in the Barnes maze were associated with decreased exploratory or increased anxiety-like behaviors, the same cohorts were evaluated at 8 weeks of age in both the elevated plus maze (EPM) and in the open field task. These tests failed to provide evidence of an anxiety-like behavior as the IL-6 Rx male and female mice spent equal amounts of time in the open arms of the EPM (**Fig.4A**), and had no differences in the total time or number of visits to the center quadrant of the open field apparatus (**Fig.4B**). Moreover, the total distance moved was assessed in the open field as a measure of exploratory behavior and revealed no differences between the IL-6 and PBS Rx mice for both males (PBS 4954±272.6 vs IL-6 5031±162.7; t(15) = 0.25, p = 0.806, unpaired t-test) and females (PBS 6103±277.2 vs IL-6 6360±177.1; t(12) = 0.78, p = 0.45, unpaired t-test).

**Fig.4.**
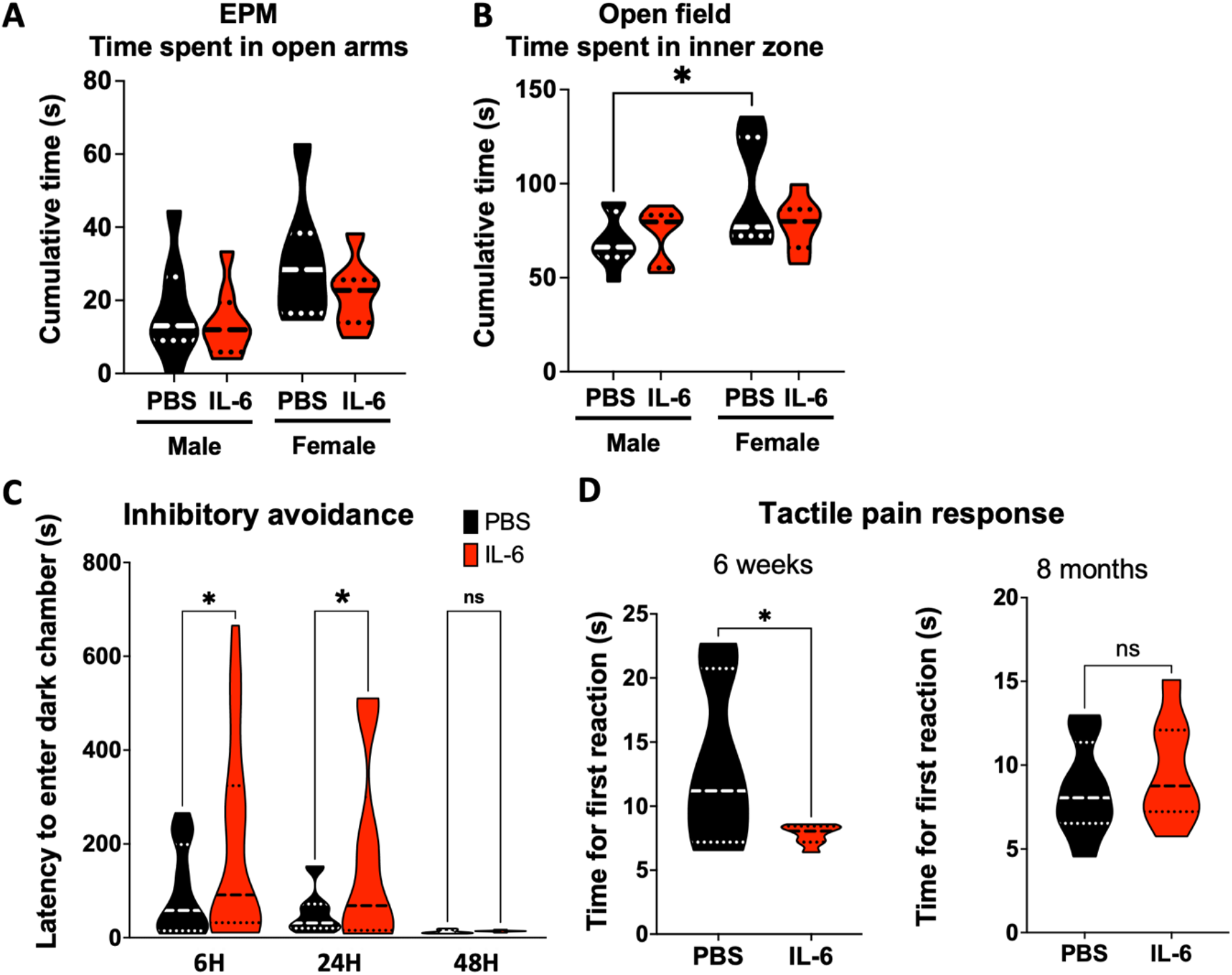
Animals exposed postnatally to IL-6 elevation have a normal response to anxiolytic stimuli but enhanced inhibitory avoidance and pain response. **A**. IL-6 Rx male and female mice were evaluated at P42 to assess anxiolytic behavior. Cumulative time (s) spent in the open arms of the Elevated Plus Maze (EPM) **B**. Cumulative time spent in the inner zone of the open field arena. Aversive memory was evaluated in the inhibitory avoidance task for the IL-6 treated and PBS control mice at P42, using the light/dark chamber. **C**. Average latency for mice to enter the dark chamber 6, 24 and 48 h after receiving the fear inducing stimulus (foot shock upon entering the dark chamber). *p<0.05 by Student’s t-test. n=19 IL-6 and 16 PBS treated mice, from 6 litters. **D**. Pain responsiveness following IL-6 treatment was evaluated in mice at 6 weeks and as adults by scoring the latency to respond to a tactile noxious heat stimulus. Latency measured as the time for any of the following responses: Licking or flicking the hind paw or jumping from the hot plate surface. *p<0.05, ns=not significant by Student’s t-test comparing IL-6 treated and PBS groups, n=8 mice per group, from 3 litters.

### IL-6 Rx mice show enhanced inhibitory avoidance learning and pain response

As the IL-6 Rx mice exhibited deficits in Barnes maze performance, which may not be associated with spatial memory, and had no signs of anxiety-like behavior in either the EPM or the open field, we evaluated mice using the inhibitory avoidance task, to assess aversive memory. At 6 weeks of age, male IL-6 Rx mice showed an enhanced inhibitory avoidance learning response at both 6h (PBS 95.17±23.19 vs IL-6 200.9±48.04) and 24h (PBS 54.59±13.10 vs IL-6 170.2±45.74) after receiving a noxious stimulus (One-way ANOVA F (5,76) = 3.975, p<0.05). At 48h post-stimulus, their performance was not different from the controls (**Fig.4C**).

Aversive memory responses can be affected by how the animals perceive a painful stimulus. Also, changes in sensory processing have been reported in models of neurodevelopmental disorders (Robertson and Baron-Cohen, 2017) and sensory processing impairments are commonly observed in individuals across the autistic spectrum (Tomchek and Dunn, 2007), including somatosensory deficits (Marco et al., 2012; Puts et al., 2014). Indeed, we had previously shown that the IL-6 Rx mice are hyposmic (Kumari et al., 2020). Therefore, we tested the IL-6 Rx male mice for their sensitivity to a tactile noxious heat stimulus. At 6 weeks, the IL-6 Rx mice had a significantly faster reaction to a mildly painful stimulus than the controls (**Fig.4D**; PBS 13.28±2.39 vs IL-6 7.8±0.27; t(13) = 2.44, p = 0.029, unpaired t-test, 3 litters). Interestingly, this increased sensitivity to a painful stimulus was no longer evident when the mice were re-tested at 8 months of age (**Fig.4D**; PBS 8.53±1 vs IL-6 9.62±1.11).

### IL-6 Rx mice have a persistent deficit in sociability

Sociability and communication deficits are often reported in mouse models for neurodevelopmental disorders and are relevant to conditions like ASD. To determine whether the IL-6 Rx would affect sociability, we evaluated IL-6 Rx 6 male mice at 6 weeks of age using the social approach test (SA). At this age, mice are normally interactive and curious, preferring to spend more time engaging with social partners. Indeed, the PBS Rx control mice spent more time investigating a naïve social partner vs an empty cage (**Fig.5A;** Empty 171.2±23.31 vs Social 294.5±28.67; t(8) = 3.34, p = 0.01. unpaired t-test). This preference for social interaction was confirmed by showing that the control mice also spent more time sniffing the social partner, measured by detecting the nose of the test animal in the sniffing zone, immediately adjacent to the partner enclosure (**Fig.5B**; Empty 96.48±13.22 vs Social 211.5±39.78; t(8) = 2.73, p = 0.026, unpaired t-test). By contrast, the IL-6 Rx mice showed no preference for interacting with another mouse vs an empty cage as measured by either time spent in the social chamber (**Fig.5A**) or in the sniffing zone (**Fig.5B,D**). In the novel social subject test (NSS), the IL-6 Rx mice again showed no preference for interacting with a naïve mouse vs. with a familiar littermate, whereas the PBS Rx mice showed a clear preference for interacting with the novel mouse (**Fig.5C,D**). We evaluated total distance traveled by the mice in the 3-chambers apparatus for all tasks to exclude confounding locomotor effects (data not shown). We further evaluated fine locomotor skill for the same animals in the horizontal ladder and found no differences in the number of foot slips between the PBS and IL-6 Rx groups (PBS 3.71±0.7 vs IL-6±0.36; t(40) = 0.06, p = 0.952, unpaired t-test). Altogether, these tests show that the IL-6 Rx mice have deficits both in engaging in social interactions and in social memory.

**Fig.5.**
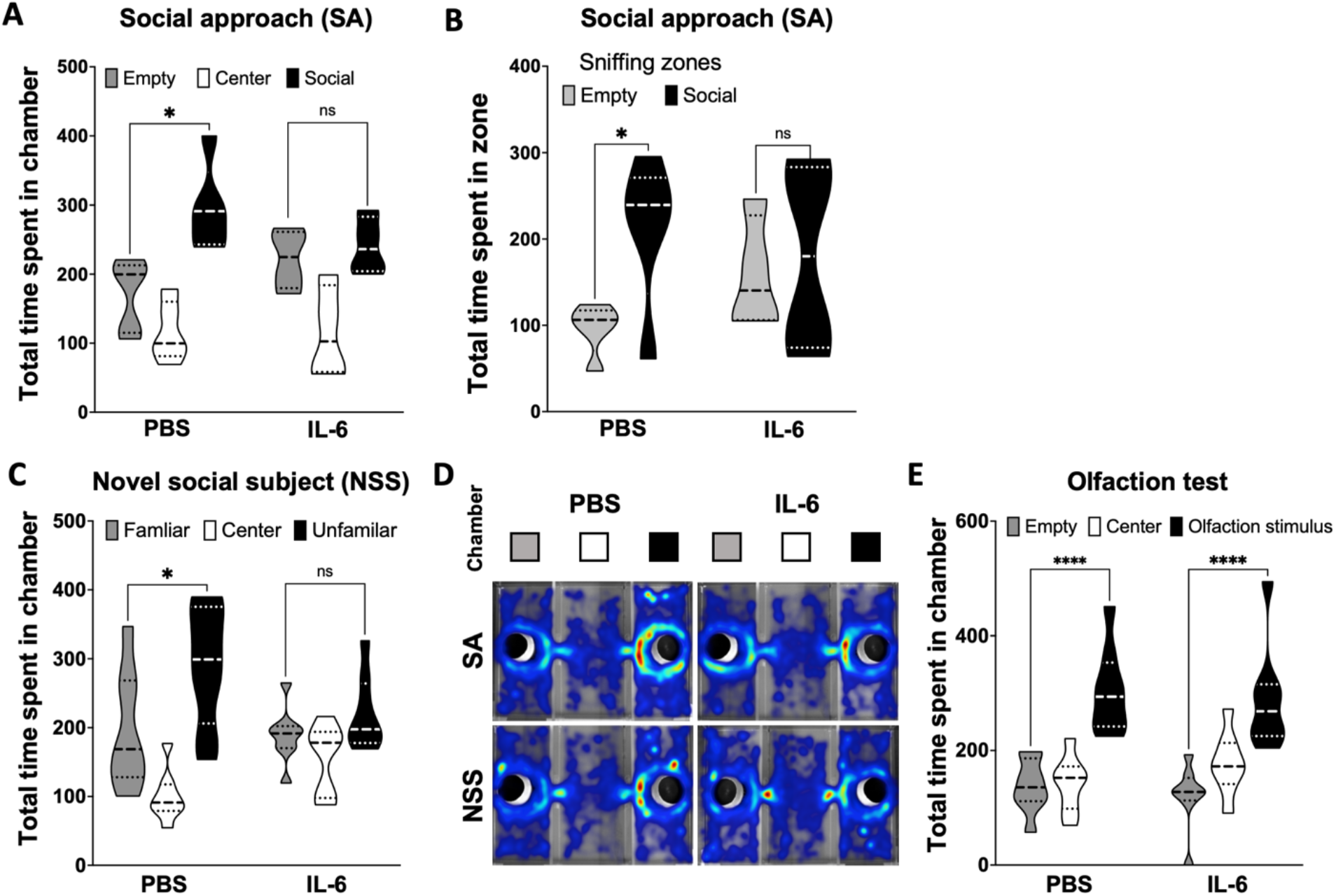
Mice exposed to IL-6 postnatally have a deficit in sociability. P42 male mice were challenged in the social approach and social novel subject tasks using an automated 3-chamber apparatus to evaluate sociability. **A**. In the Social approach test (SA), total time spent in the non-social (empty), center and social (naïve social partner) chambers was recorded via automated video tracking for IL-6 treated and PBS controls. **B**. Cumulative time in the sniffing zone (15 cm radius around partner enclosure). **C**. Mice were challenged in the novel social subject task (NSS), where they were introduced to known (familiar chamber) and socially naïve (unfamiliar chamber) partners. Time spent in each of the three chambers was recorded **D**. Heatmaps for the group average of time spent in each chamber for PBS and IL-6 animals in the Social approach (SA) and Novel Social Partner (NSS) tasks. **E**. Olfaction stimulus test in the 3-chambers. Total time spent in an empty chamber or in a chamber with a strong olfactory stimulus (mouse urine). n=8 mice per group, from 3 litters. *p<0.05 by Student’s t-test comparing IL-6 treated and PBS groups.

Since mice use their sense of olfaction to detect social partners, and as we had previously shown that the IL-6 Rx mice are hyposmic, we hypothesized that SA and NSS performance could have been affected by altered olfactory function. To test this hypothesis, we performed the 3-chamber test again, but replaced the live social partner with filter paper soaked in mouse urine and covered both the cages so that the test mouse could not easily tell that there wasn’t a mouse in one of the cages. Using this variation of the test, both the PBS Rx and the IL-6 Rx mice showed a clear preference for investigating the chamber with the olfactory stimulus, showing that the deficits observed in the SA and NSS cannot be attributed simply to a defect in olfaction (**Fig.5E**; 27.21±3.3 vs 79.61±6.57; t(10) = 7.13, p<0.0001, unpaired t-test).

### Animals transiently exposed to IL-6 postnatally have a persistent increase in repetitive behaviors

Exacerbated self-grooming behavior in mice is analogous to the repetitive, stereotyped behaviors often observed in individuals with ASD and with certain other neurodevelopmental disorders (Tracy et al., 1996; Howlin et al., 2004; Malkova et al., 2012); therefore, we evaluated grooming behavior at 3 months of age. This analysis revealed that the male IL-6 Rx mice spent more time grooming themselves than the PBS Rx mice (**Fig.6A;** PBS 35.54±4.94 vs. IL-6 58.82±8.52; t(15) = 2.28, p = 0.037, unpaired t-test), indicating a persistent increase in repetitive behavior. Moreover, age-matched female littermates had a similar increase in time spent self-grooming (**Fig.6A;** PBS 25±3.5 vs. IL-6 63.05±9.58; t(11) = 3.97, unpaired t-test). Therefore, a transient postnatal exposure to IL-6 appears to cause a sex-independent increase in repetitive behaviors in mice.

**Fig.6.**
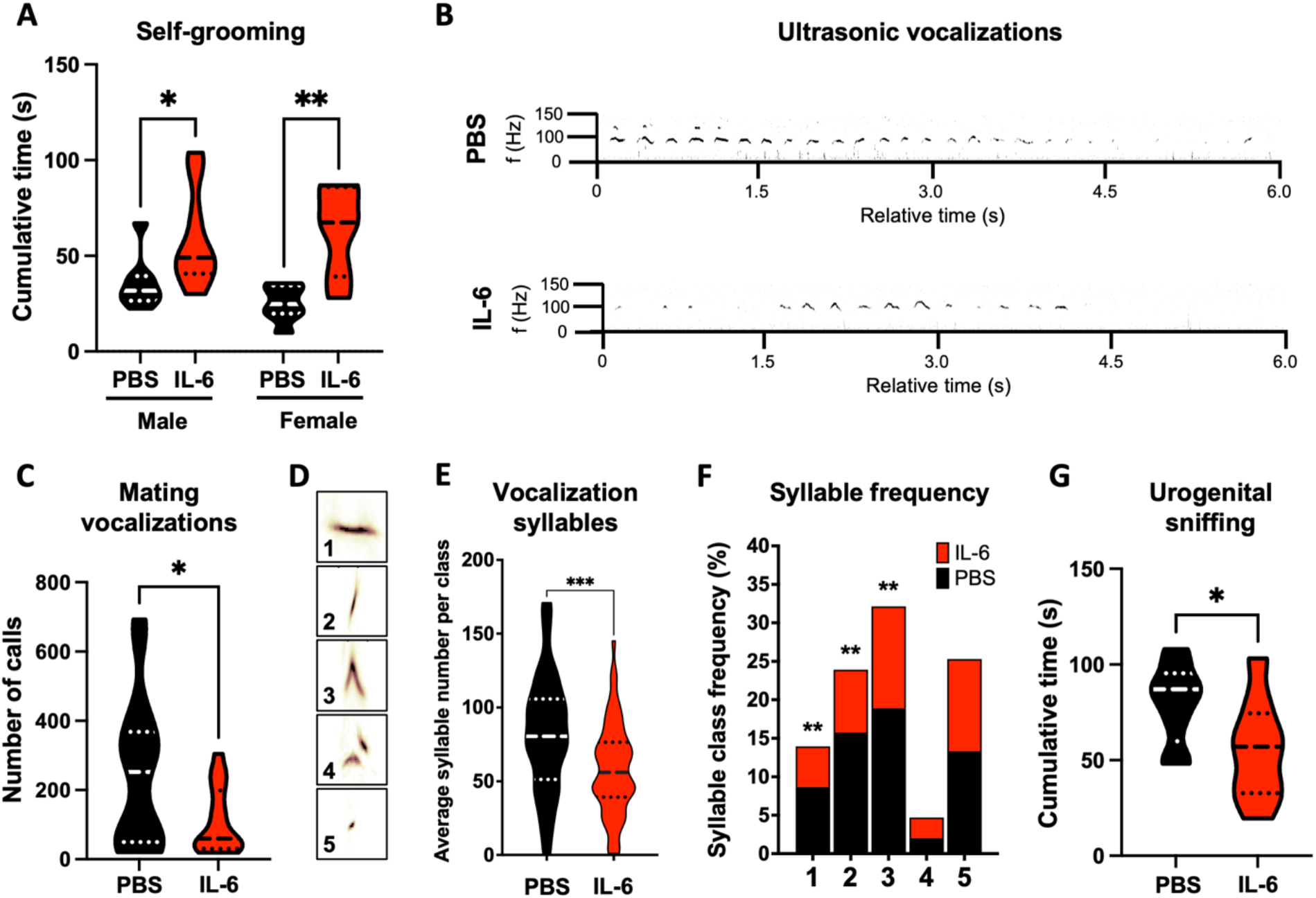
Animals exposed to IL-6 postnatally have a persistent increase in repetitive behaviors and aberrant mating interactions. Self-grooming was evaluated as a measure of repetitive behaviors for IL-6 injected mice at 3 months of age. **A**. Cumulative duration (s) of grooming behaviors in adult male and female mice in isolated trials. * p<0.05, ** p<0.01 by Student’s t-test, n=8 mice per group from 3 litters. **B**. Representative track of recorded ultrasonic vocalizations. **C**. Number of ultrasonic vocalization (USV) calls recorded during mating interactions. * p<0.05 by Student’s t-test, n=8 mice per group from 3 litters **D**. Syllables from USVs were classified into 5 classes using the MUPET software. **E**. Average number of vocalization syllables from PBS (black) and IL-6 treated (red) mice per syllable class. *** p<0.001 by Student’s t-test, n=12 mice per group from 3 litters **F**. The relative frequency of each syllable for each of the 5 classes was quantified. ** p<0.01, by Student’s t-test, n=12 mice per group from 3 litters. **G**. Cumulative time spent engaging in urogenital sniffing by the male partner during mating interactions. *p<0.05 by Student’s t-test. n=12 mice for each group.

### Aberrant social behaviors are persistent in adult mice exposed to IL-6 postnatally

Male mice display characteristic behaviors during courtship in response to female odors, which include producing ultrasonic vocalizations (USV) (Nyby et al., 1977). Therefore, we recorded and evaluated USVs when the males were in the presence of a sexually receptive female. Interestingly, IL-6 Rx produced significantly fewer USV calls during these encounters, compared to PBS Rx littermates (**Fig 6B,C**; PBS 246,7±51.63 vs. IL-6 110.07±23.76; t(31) = 2.34, p = 0.026, unpaired t-test, 6 litters). Their USVs had normal physical characteristics (duration, frequency, amplitude, bandwidth, syllable morphology; data not shown), indicating a social deficit rather than a physical inability to produce calls. However, when the syllables were sorted into 5 classes, the IL-6 Rx animals produced fewer syllables on average than the PBS Rx mice (**Fig.6D**). Also, comparing the syllable classes individually revealed that IL-6 Rx mice produced fewer calls with flat (class 1), frequency-up (class 2) and chevron (class 3) syllables (**Fig.6E,F**; class 1: PBS 73.1±9.13 vs. IL-6 44.6±10.6, t(9) = 3.98, p = 0.003; class 2: PBS 100±12.36 vs. IL-6 62.55±7.08, t(10) = 4.17, p = 0.002; class 3: PBS 88.67±7.58 vs. IL-6 61.94±6.46, t(17) = 6.08, p<0.0001; paired t-test). There were no changes in those syllables with 2-frequency step (class 4) or with short punctate syllables (class 5) **(Fig.6F)** (Heckman et al., 2016).

When courtship behaviors were scored, there was a trend for the IL-6 Rx mice to have an increased latency before their first attempt to mount a female and a similar trend for fewer total attempts to mount the females (data not shown). While these trends did not reach statistical significance, the IL-6 Rx mice spent significantly less time sniffing the urogenital region of the females (**Fig.6G**; PBS 80.01±6.92 vs. IL-6 55.38±8.78; t(16) = 2.21, p = 0.0424, unpaired t-test), reinforcing the hypothesis of mating deficits in these mice.

## Discussion

Epidemiologic studies and rodent models have linked increased circulating levels of IL-6 to cognitive and behavioral abnormalities. Our previous studies, as well as work from others, have shown that elevating systemic levels of IL-6 changes the proliferation and subsequently the composition of progenitors present in the secondary germinal matrices of the brain, namely, the SVZ and SGZ (Monje et al., 2003; Covey et al., 2011; Wei et al., 2011; Kumari et al., 2020). Notably, IL-6 Rx decreased the proliferation of two types of glial restricted progenitors in the SVZ while increasing the proliferation of a multipotent progenitor. Here, we used Nestin-CreERT2/Rosa26-TdTomato reporter mice to follow the progeny of the NSCs and found that perinatal IL-6 Rx reduced the number of protoplasmic astrocytes in the amygdala. This finding was un-expected as previous studies had reported that IL-6 increased the differentiation of embryonic NSCs and the specification of neocortical progenitors into astrocytes (Bonni et al., 1997; Nakanishi et al., 2007). However, our findings are consistent with the reported reduction in the number of layer II and III astrocytes in the frontal cortex of autism patients (Falcone et al., 2021). Since astrocytes support synaptogenesis and synaptic function, the decrease in astrocytes subsequent to IL-6 Rx may contribute to increases or decreases in synapse numbers (Allen and Eroglu, 2017). For example, injecting newborn astrocytes into the adult brain is sufficient to recover cortical deficits in synaptic plasticity in the ocular dominance columns (Muller and Best, 1989). Furthermore, ASD-derived astrocytes interfered with the development and function of control neurons, while astrocytes from control patients rescued the synaptogenesis defects seen with ASD-derived neurons. Furthermore, blocking IL-6 secretion from the ASD-derived astrocytes improved synaptogenesis (Russo et al., 2018). Similarly, astrocytes from the Rett syndrome (RTT) mouse, that lack methyl-CpG-binding protein 2 (MeCP2), failed to support normal morphological maturation and function of wild type neurons in co-cultures (Ballas et al., 2009). Similarly, human MeCP2-deficient astrocytes differentiated from RTT induced pluripotent stem cells negatively affect the morphology and electrophysiological function of GABAergic interneurons (Williams et al., 2014). Re-expressing MeCP2 preferentially in astrocytes of globally MeCP2-deficient mice, rescued locomotion and anxiety levels and greatly prolonged lifespan compared to globally null mice. Furthermore, replacing MeCP2 in the mutant astrocytes restored dendritic morphology and glutamate metabolism (Lioy et al., 2011).

Altogether these data support the hypothesis that impairing astroglial development in the forebrain will disturb synaptogenesis with long lasting behavior deficits. Supporting this hypothesis, a prospective study that analyzed a large cohort of ASD patients found that the largest subset of ASD candidate genes were associated with synaptic transmission and glial cell differentiation and they were differentially expressed in the corpus callosum of ASD patients (Li et al., 2014). While studies have shown that the white matter is abnormal in psychiatric patients as discussed below, the proposition that stunted astrocyte development contributes to the dysgenesis of the forebrain and to the origin of neurodevelopmental psychiatric disorders has received less support.

An intriguing possibility is that the IL-6 induced reduction in astrocytes might be due to increased microglia phagocytosis. VanRyzin et. al., (2019) showed that in early postnatal mouse CNS development microglia preferentially engulf newborn astrocytes. Blocking phagocytosis in the amygdala with a CD11b antibody was sufficient to increase the number of mature astrocytes, that in turn, restored juvenile rough-and-tumble play behavior (VanRyzin et al., 2019). While a possibility, it is not likely that phagocytosis is contributing to the astrocyte deficiency reported here since microglial phagocytosis peaks at P0, which is one day before we labelled the NCSs that would eventually produce TdTomato+ astrocytes in these studies.

Reduced numbers of oligodendrocytes are seen in postmortem brains from ASD patients older than 20 years of age (Morgan et al., 2014) and white matter alterations are seen in patients with Schizophrenia and ASD (Bartzokis, 2012). Therefore, we evaluated gliogenesis in the white matter tracts and found that the IL-6 Rx mice had fewer oligodendrocytes in both the body and the splenium of the corpus callosum. These results were not predictable as *in vitro* studies had shown that IL-6 promoted the maturation of oligodendrocyte progenitor cells (OPC) into oligodendrocytes (Valerio et al., 2002). Furthermore, an analysis of the frontal lobe of BTBR mice had precocious myelination and increased white matter volume (Khanbabaei et al., 2019). Moreover, neurospheres derived from embryonic NSCs treated with IL-6 differentiate into OPCs with increased myelinating capacity (Zhang et al., 2006). However, consistent with our results an analysis of 5 different mouse ASD models showed disruptions in genes that regulated oligodendrocyte biology (Phan et al., 2020).

Nestin-CreER(T2)/R26R-YFP reporter mice have been used to show that the progeny of adult nestin-expressing stem cells in the adult SVZ and the SGZ produce neuroblasts that differentiate into olfactory bulb interneurons and hippocampal dentate gyrus excitatory granule neurons (Lagace et al., 2007). In our previous cell fate mapping studies, we showed that perinatal IL-6 Rx did not affect olfactory bulb interneuron genesis (Kumari et al., 2020). By contrast, here we show that IL-6 Rx significantly reduced hippocampal granule cell layer neurogenesis. Similarly, transgenically increasing IL-6 production from astrocytes decreased neurogenesis by 63% in the dentate gyrus (Vallieres et al., 2002). We observed a similar repression of hippocampal neurogenesis with IL-1β Rx using this same administration paradigm (Veerasammy et al., 2020b). These data have important ramifications for human brain development and function. While hippocampal neurogenesis continues across the lifespan in rodents, studies have shown that a proliferative germinal zone disappears from the human hippocampus during childhood and that adult hippocampal neurogenesis is either extremely rare or non-existent (Taylor et al., 2013; Sorrells et al., 2018); therefore, perturbations to human hippocampal development will likely have persistent consequences that will contribute to deficits in learning and memory, social memory, contextual fear and emotional control (Fanselow and Dong, 2010).

To establish how the increased levels of IL-6 during this late period would affect behavior we subjected the mice to tasks pertinent to core symptoms of neurodevelopmental disorders. To assess anxiety both male and female mice were evaluated using the elevated plus maze and the open field test but neither test showed any changes in exploration or anxiety-like behavior. These data are in contrast with those reported using MIA models that administered LPS or poly(I:C) mid-gestation. Increased anxiety-like behaviors have been seen in adult offspring of rat dams injected with LPS from gestational day 7 to birth (Chamera et al., 2020), in offspring of mouse dams injected with poly(I:C) at E12.5 (Hsiao et al., 2012), and in rats injected with either LPS or poly(I:C) at midgestation injections (Talukdar et al., 2020). Notably, directly activating maternal T-cells during pregnancy using Staphylococcal Enterotoxin A and B induced anxiety-like behavior in the offspring with males exhibiting higher anxiety (Glass et al., 2019). Interestingly, when Carlezon et al., used a “dual-hit” immune activation model that combined Poly(I:C)-induced mid-gestational MIA followed by LPS administration on postnatal day 9, there was a male specific increase in anxiety-like behavior as assessed by the open field test (Carlezon et al., 2019). The data suggest that the propensity for increased anxiety can be attributed to the effects of inflammation on the progenitors or their progeny of the primary germinal zones during fetal development and not to altered secondary germinal zone neurogenesis.

Freezing behavior is often associated with fear responses; therefore, we explored this phenotype in the IL-6 Rx mice using the inhibitory avoidance task. Juvenile male IL-6 Rx showed an enhanced inhibitory avoidance memory. One caveat here, however, is that the IL-6 Rx mice may have been more sensitive to the foot-shock as they responded more strongly to a noxious tactile stimulus when tested at 6 weeks with the hot-plate test. The circuitry underlying aversive responses is complex with multiple brain regions involved that include the ventral hippocampus, ventral striatum, amygdala and prefrontal cortex. Silencing the activity of ventral hippocampal neurons increases stress susceptibility and that increasing hippocampal neurogenesis makes mice more stress resistant (Anacker et al., 2018), reminiscent of our data on hippocampal neurogenesis. However, the increase in inhibitory avoidance memory may be linked to changes in the prefrontal cortex (PFC) and amygdala. The infralimbic region of the PFC is strongly connected to both the hippocampus and amygdala and regulates fear, anxiety and stress in many paradigms including fear conditioning (Yizhar et al., 2011). Stress can increase the consolidation of aversive memories via glucocorticoid activity in the amygdala (Roozendaal et al., 2008). The reduced number of astrocytes in the amygdala elicited by IL-6 Rx might affect this circuit by impairing astrocytic glutamate uptake.

Male mice produce characteristic USV, which are most prominent during their initial encounters with females and correlate well with the level of male sexual arousal (White et al., 1998). We found that the adult male IL-6 Rx mice emitted significantly fewer USVs when paired with receptive females. Moreover, this effect correlated with a reduction in partner investigatory behavior (urogenital sniffing), indicating that the IL-6 Rx males had reduced sexual interest. However, we cannot fully exclude that the decreased urogenital sniffing wasn’t affected by their mild hyposmia. Communication deficits during mating have been previously reported in MIA models, notably, in the study by Malkova et al., 2012, where the male offspring from MIA dams emitted significantly fewer USVs in response to encounters with females, and they also displayed reduced scent marking in response to female urine (Malkova et al., 2012). Notably, several genetic mouse models relevant to autism (BTBR, NL-3-knockout [KO], NL-4 KO and Shank1 KO) show a similar deficit in mating elicited USVs (Jamain et al., 2008; Radyushkin et al., 2009; Scattoni et al., 2011; Wohr et al., 2011).

We further investigated the social interactions of the mice using the 3-chamber apparatus where the IL-6 Rx mice displayed a lack of social preference at 6 weeks, an age when mice are normally interactive and curious. The IL-6 Rx mice also showed reduced preference for a novel mouse vs. a familiar mouse in the novel social partner task suggesting that the IL-6 Rx mice have problems with social memory. Importantly, since we previously reported that exposing mice postnatally to IL-6 induced mild hyposmia (Kumari et al., 2016), we challenged the IL-6 injected mice in the 3-chamber apparatus with mouse urine soaked filter paper vs. clean filter paper and showed that like the PBS Rx mice, the IL-6 Rx mice had a normal preference for investigating the chamber that smelled of mouse urine. These data support the conclusion that their failure to investigate the mouse in the social approach test cannot be attributed to a defect in olfaction.

The social deficits we observed in the IL-6 Rx male mice align well with previous observations in MIA rodent models. The offspring of rats born to dams exposed to either LPS or poly(I:C) midgestation show decreased social approach and social memory when tested in the 3-chamber test at both peri-adolescence (6 weeks) and as early adults (10 weeks) (Talukdar et al., 2020). Similarly, 10 week old C57BL/6J mice born from poly(I:C) (Malkova et al., 2012) and Staphylococcal Enterotoxin (Glass et al., 2019) treated dams displayed lack of social preference in the 3-chamber social approach task. Furthermore, in the “dual-hit” model introduced by Carlezon et al., 2019, social approach was significantly reduced in 8-week-old male mice but unchanged in the females, suggesting male-specific deficits in this behavioral domain (Carlezon et al., 2019). Importantly, blocking the increase in circulating IL-6 in MIA models improved social deficits in rats and mice. In an LPS rat MIA model, the postnatal administration of the anti-inflammatory pioglitazone reduced plasma levels of IL-6 and was sufficient to improve LPS induced deficits in juvenile social play (Kirsten et al., 2018). Similarly, neutralizing IL-6 with a function blocking antibody in the offspring of poly(I:C) treated mice prevented latent inhibition and pre-pulse inhibition. Moreover, a single dose of anti-IL-6 antibody was sufficient to prevent the social approach deficits caused by poly(I:C) administration (Smith et al., 2007).

Repetitive and stereotyped behaviors are core symptoms in ASD and they are observed in Schizophrenia. Furthermore, serum levels of IL-6 have been directly associated with stereotypical behaviors in children with ASD (Ashwood et al., 2011). Genetic mouse models for these psychiatric disorders often exhibit increases in characteristic repetitive behaviors that are interpreted as reflective of patients’ stereotypical movements. For instance, mice with mutations in the neuroligin-3 gene, which is associated with ASD in humans, have increased stamina on the rotarod test, which was interpreted as an increase in repetitive behavior (Rothwell et al., 2014). Increase in rotarod performance was also reported for male mice born from LPS treated dams (Carlezon et al., 2019). However, increased self-grooming is commonly used as a mouse analog for repetitive/stereotyped behaviors and this behavior has been observed in multiple genetic mouse strains relevant to ASD (BTBR, Fmr1, MeCP2 and CNTNAP2 KO mice) (McFarlane et al., 2008; McNaughton et al., 2008; Chao et al., 2010; Penagarikano et al., 2011). Increases in self-grooming also have been observed in the offspring of poly(I:C) injected dams (Malkova et al., 2012). Here we have shown that both male and female mice spend significantly more time self-grooming compared to PBS Rx controls. To our knowledge, this is the first report including a sex comparison for this task in a perinatal inflammation animal model. Altogether, these assays demonstrate that the perinatal IL-6 Rx mice exhibit many negative signs seen in neurodevelopmental disorders such as ASD and schizophrenia.

Overall, the data presented here indicate that transient neonatal exposure to IL-6 reprograms SVZ and SGZ-derived neural stem cells and progenitors, altering the fates and distribution of their progeny, which subsequently alters the cellular composition of several late developing brain structures. Whereas previous studies using antenatal administrations of LPS, Poly(I:C) or cytokines have shown that those mice have behavioral deficits, their administration paradigms would have affected the NSCs and progenitors of both the primary and the secondary germinal zones; therefore, precluding any conclusions as to which progenitors were affected and whether there is a critical period for exposure. Our data show that perturbations that only affect the stem cells and progenitors in the secondary germinal matrices are sufficient to elicit long-lasting changes across a range of cardinal behaviors related to neurodevelopmental disorders.

## Acknowledgments

Research reported in this publication was supported by the Governor’s Council on Autism, CAUT22AFP009 awarded to FJV, CAUT17BSP010 awarded to SWL and from a grant from the National Institutes of Health R21 NS107772 awarded to SWL.

